# A CO_2_-limitation-induced cytosolic repressor enables shutdown of the algal CO_2_-concentrating mechanism

**DOI:** 10.64898/2026.09.01.748463

**Authors:** Shizuka Miichi, Daisuke Shimamura, Junko Yasuda, Ryutaro Tokutsu, Takashi Yamano

**Author notes:** **Corresponding author:** Takashi Yamano, Graduate School of Biostudies, Kyoto University, Kyoto, 606−8502, Japan. **Author contributions:** S.M. and T.Y. designed research; S.M., D.S., and J.Y. performed research; R.T. and T.Y. contributed new reagents/analytic tools; S.M., J.Y., and T.Y. analyzed data; and S.M., D.S., and T.Y. wrote the paper.

## Abstract

Aquatic photosynthetic organisms face limited CO_2_ availability because CO_2_ diffuses slowly in water and most dissolved inorganic carbon (Ci) exists as HCO_3_^−^ at physiological pH. To overcome this limitation, aquatic photoautotrophs operate CO_2_-concentrating mechanisms (CCMs) that elevate CO_2_ around Rubisco and sustain carbon fixation. Because CCM operation consumes energy, it must be suppressed when CO_2_ becomes abundant, but how this shutdown occurs remains poorly understood. In *Chlamydomonas reinhardtii*, the nuclear protein CBP1 was identified as a CCM repressor, but its loss causes only partial derepression under high CO_2_, indicating that an additional mechanism is required for complete shutdown. Here, we identify High-Affinity CCM Repressor 1 (HCR1), a cytosolic protein related to CBP1, as a second repressor. Under high CO_2_, *hcr1* mutants retained high affinity for Ci and derepressed CCM and photoacclimation genes. Combined disruption of *HCR1* and *CBP1* further increased Ci affinity, approaching that of wild-type cells with a fully induced CCM under CO_2_ limitation, and promoted the accumulation of Ci transporters. *HCR1* loss also prevented redistribution of the chloroplast regulator CAS away from the pyrenoid and was accompanied by retention of a pyrenoid starch sheath. In contrast, LCIB, a chloroplast CO_2_-recapture protein, relocated normally. Unexpectedly, HCR1 accumulated during CO_2_ limitation and declined after transfer to high CO_2_. These results show that CCM shutdown is an active transition rather than the passive reversal of induction. We propose that CBP1 restrains CCM1-dependent transcription, while HCR1 is preloaded during CO_2_ limitation to terminate the CAS-associated, starch-sheathed, high-affinity state when CO_2_ becomes replete.

**Significance Statement:** Aquatic photosynthetic organisms often expend energy to concentrate CO_2_ around Rubisco, the enzyme that fixes CO_2_ into organic carbon. Research has emphasized how this system is activated under CO_2_ limitation, while its shutdown has usually been viewed as a simple reversal. We show that shutdown is instead an active process that coordinates gene repression, chloroplast signaling, and remodeling of the pyrenoid, the compartment where Rubisco and CO_2_ are concentrated. Unexpectedly, part of the shutdown machinery accumulates while CO_2_ remains limiting, suggesting that cells prepare the off-switch before conditions improve. This finding suggests a general strategy for reversible environmental adaptation and offers a framework for engineering photosynthesis that captures carbon efficiently without wasting energy as CO_2_ availability changes in fluctuating environments.

## Introduction

Ribulose-1,5-bisphosphate carboxylase/oxygenase (Rubisco) catalyzes the entry of CO_2_ into the Calvin–Benson cycle, but its slow catalytic turnover and competing oxygenation reaction constrain photosynthetic carbon fixation. These limitations are especially acute in aquatic environments, where CO_2_ diffuses much more slowly than in air and, at circumneutral pH, most dissolved inorganic carbon (Ci) is present as HCO_3_^−^ rather than CO_2_. Consequently, the CO_2_ concentration available to Rubisco is frequently below saturation. Many aquatic photoautotrophs compensate for this limitation by operating a CO_2_-concentrating mechanism (CCM) that actively accumulates Ci and elevates the CO_2_ concentration around Rubisco (1–3).

In the model green alga *Chlamydomonas reinhardtii*, the CCM is organized around the pyrenoid, a Rubisco-rich compartment within the chloroplast (4, 5). The pyrenoid matrix assembles through phase separation of Rubisco with the linker protein EPYC1 (6–8). It is traversed by thylakoid-derived tubules and, under CO_2_-limiting conditions, is surrounded by a starch sheath that contributes to CO_2_ retention (9). HCO_3_^−^ uptake is mediated by HLA3 at the plasma membrane and LCIA at the chloroplast envelope (10–14). HCO_3_^−^ is subsequently delivered to the thylakoid lumen through the bestrophin-like proteins BST1–3, where the lumenal carbonic anhydrase CAH3 generates CO_2_ close to the pyrenoid matrix (15, 16). Under less severe CO_2_ limitation, the periplasmic carbonic anhydrase CAH1 and the plasma-membrane protein LCI1 support an additional CO_2_-uptake route (17, 18).

The *Chlamydomonas* CCM is not a binary on–off system. Instead, distinct low-CO_2_ (LC) and very-low-CO_2_ (VLC) acclimation states are engaged as dissolved CO_2_ declines (19, 20). LC-acclimated cells rely predominantly on a CO_2_ uptake pathway involving CAH1 and LCI1, whereas VLC acclimation additionally engages the high-affinity HCO_3_^−^ uptake system involving HLA3 and LCIA. The two states are also distinguished by chloroplast remodeling. The LCIB–LCIC complex remains dispersed throughout the chloroplast under HC and LC conditions but relocalizes to the pyrenoid periphery when dissolved CO_2_ falls below approximately 7 µM, marking entry into the VLC state (20, 21). VLC acclimation is accompanied by enrichment of CAS along pyrenoid tubules and development of a conspicuous pyrenoid starch sheath (9, 22). Because photosynthetic CO_2_ consumption progressively lowers dissolved CO_2_ in liquid cultures, aeration with ambient air containing approximately 0.04% CO_2_ drives cells into the VLC state rather than maintaining a generic LC condition (20, 23).

Transcriptional induction of the CCM is controlled primarily by CCM1/CIA5, hereafter CCM1, a nuclear zinc-finger regulator required for the expression of many CO_2_-limitation-responsive genes (24–27). CCM1 acts in part through downstream regulators such as the Myb transcription factor LCR1 and can directly bind a GC-rich promoter motif (28, 29). A second regulatory layer originates in the chloroplast. The thylakoid-associated Ca^2+^-binding protein CAS redistributes from thylakoid membranes throughout the chloroplast under HC to pyrenoid tubules under VLC and is required to maintain the expression of a subset of nuclear CCM genes, including the Ci-transporter genes *HLA3* and *LCIA* (22, 30, 31). Disruption of pyrenoid architecture can impair both CAS localization and CAS-dependent nuclear gene expression, indicating that the pyrenoid functions not only as a carbon-fixation compartment but also as a regulatory platform for chloroplast-to-nucleus signaling (31).

Because Ci transport, thylakoid lumen acidification, and localized CO_2_ generation require substantial photosynthetic energy, maintaining the CCM when CO_2_ is abundant imposes an avoidable energetic cost (23, 32, 33). When CO_2_ becomes abundant and can reach Rubisco predominantly by diffusion, cells must therefore rapidly suppress Ci transport, CCM-related gene expression, and the chloroplast architecture associated with the high-affinity state. Nevertheless, whereas the pathways that activate the CCM under CO_2_ limitation have been characterized in considerable detail, the mechanisms that actively terminate these responses after CO_2_ becomes replete remain poorly understood.

A first step toward closing this gap came with the identification of CCM1-binding protein 1 (CBP1), the first molecular repressor of the green-algal CCM (34). CBP1 is a nuclear COG0523-family protein that interacts with CCM1 and restrains CCM1-dependent transcription under HC conditions. However, CBP1 accounts for only part of CCM shutdown. Under HC, *cbp1* cells derepress only a subset of CCM1-dependent genes and retain an intermediate Ci-affinity phenotype rather than acquiring the full high-affinity state of VLC-acclimated cells. In light of the distinct LC and VLC acclimation programs, this partial phenotype suggested that an additional repressive mechanism prevents inappropriate engagement or persistence of the VLC-specific high-affinity CCM. The identity of this mechanism and its relationship to chloroplast remodeling remained unknown.

Here, we identify *Cre16.g685050*, previously designated *LCI15*, as High-Affinity CCM Repressor 1 (HCR1), a second negative regulator of the *Chlamydomonas* CCM. Genetic and physiological analyses show that HCR1 and CBP1 act as complementary repressive checkpoints, with HCR1 preferentially constraining the high-affinity CCM state under HC conditions. HCR1 is also required for CO_2_-responsive redistribution of CAS and remodeling of pyrenoid-associated starch. These findings establish CCM shutdown as an active, multilayered process that coordinates nuclear transcriptional repression with chloroplast retrograde signaling and pyrenoid architecture.

## Results

### Identification of HCR1, a VLC-induced protein containing CobW_C and WW domains

To identify a CBP1-related factor that might contribute to repression of the CCM, we surveyed the *Chlamydomonas reinhardtii* genome for COG0523/CobW-related proteins. A neighbor-joining analysis of 14 proteins recovered a four-member clade containing CBP1 (Cre16.g684650), Cre16.g685050, Cre16.g685100, and Cre16.g685000 with 99% bootstrap support. Within this clade, Cre16.g685050 grouped with Cre16.g685100 and Cre16.g685000 with 95% support, whereas CBP1 formed the sister branch (Fig. 1A). CBP1 and Cre16.g685050 were nevertheless the only proteins in this set containing a WW domain. We therefore focused on Cre16.g685050, which lacks the N-terminal CobW domain of CBP1 but retains the C-terminal CobW_C and WW domains (Fig. 1A). In C9 cells, its mean transcript abundance increased from approximately 3.6 log_2_(CPM + 1) under HC to 7.8 under VLC, corresponding to roughly 20-fold higher mean CPM under VLC (Fig. 1A). *Cre16.g685050*, *Cre16.g685100*, and *Cre16.g685000* form a compact cluster on chromosome 16 and are transcribed in the same direction, a configuration consistent with local tandem duplication (Fig. 1B). *Cre16.g685050* was originally designated low-CO_2_-inducible 15 (*LCI15*) because of its induction by CO_2_ limitation. Based on the mutant phenotype established below, we hereafter refer to LCI15 as High-Affinity CCM Repressor 1 (HCR1).

**Fig. 1.**
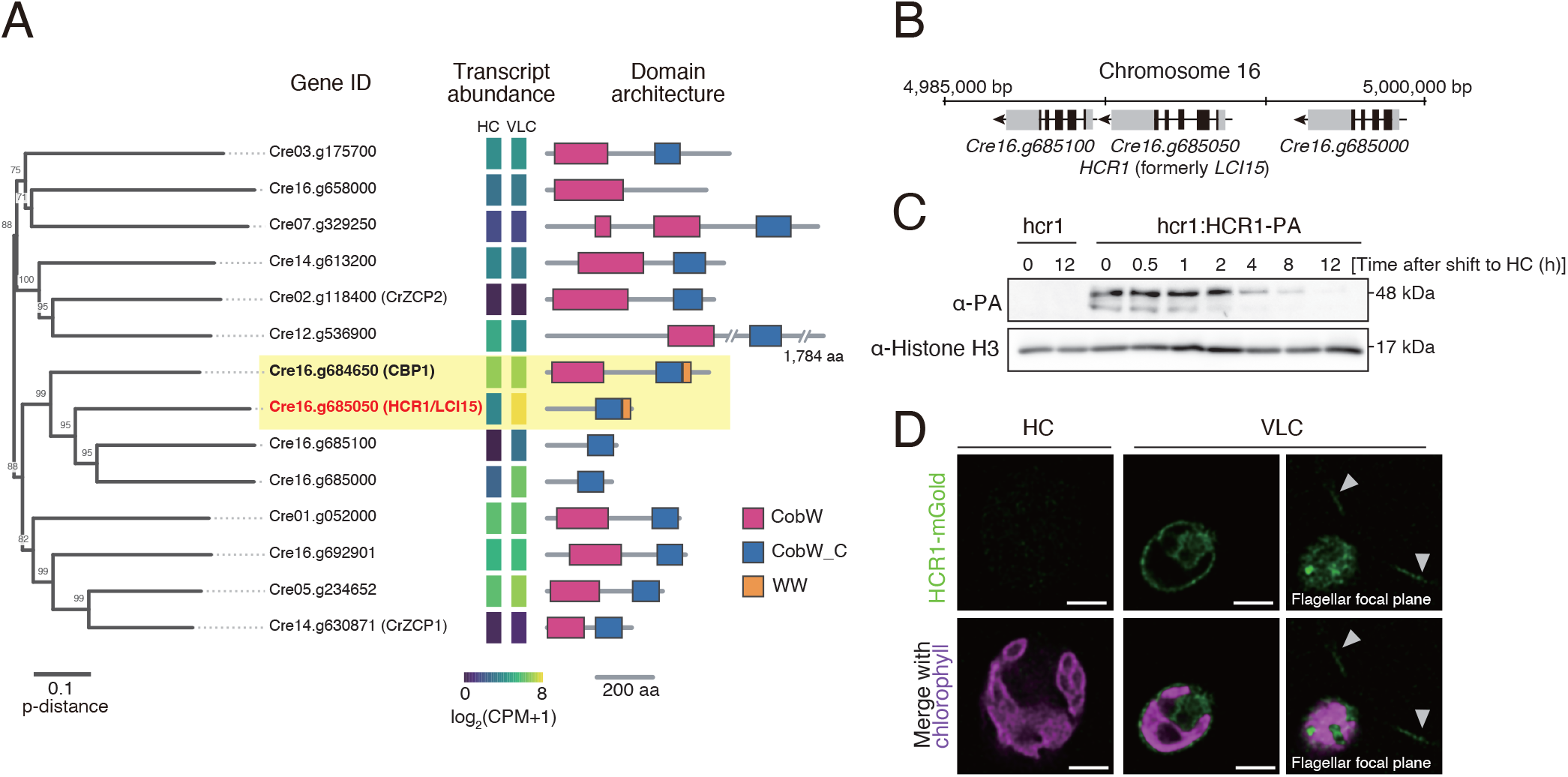
Identification, CO_2_-responsive abundance, and localization of HCR1. **(A)** Neighbor-joining tree of the 14 *Chlamydomonas reinhardtii* COG0523/CobW-related proteins, shown with mean transcript abundance in C9 under HC and VLC (*n* = 2 biological replicates) and predicted domain architectures. Node labels indicate bootstrap support ≥ 70% from 1,000 replicates. Scale bars represent a p-distance of 0.1 and 200 aa, respectively. CobW, magenta; CobW_C, blue; WW, orange. Yellow shading marks CBP1 and HCR1; HCR1 is labeled in red. The interrupted backbone denotes the 1,784-aa Cre12.g536900 protein. **(B)** Genomic organization of *Cre16.g685100*, *HCR1* (formerly *LCI15*; *Cre16.g685050*), and *Cre16.g685000* on chromosome 16. Black boxes, coding exons; gray boxes, untranslated regions; lines, introns; arrowheads, transcriptional direction. **(C)** Immunoblot of HCR1-PA during transfer of *hcr1*:*HCR1-PA* cells from VLC (0.04% CO_2_) to HC (5% CO_2_). Untagged *hcr1* cells served as negative controls, and histone H3 was the loading control. The 48-kDa marker position is indicated. **(D)** Confocal images of HCR1-mGold-PA expressed from the native *HCR1* promoter in the *hcr1* background under HC and VLC. HCR1-mGold fluorescence is green and chlorophyll autofluorescence is magenta. The right VLC column shows a flagellar focal plane; arrowheads indicate flagella. The HC and corresponding VLC cell-body images were acquired and displayed using identical settings. Scale bars, 5 µm.

To examine HCR1 protein abundance, we introduced a native-promoter HCR1-PA complementation construct into the *hcr1* background, generating a line hereafter designated *hcr1:HCR1-PA*, and monitored HCR1-PA abundance during a shift from VLC to HC. A PA-reactive band migrating near the 48-kDa marker was detected in *hcr1:HCR1-PA* but not in the untagged *hcr1* control (Fig. 1C). The signal remained strong from 0 to 1 h after transfer to HC, decreased markedly between 2 and 4 h, and was barely detectable by 8 to 12 h. Thus, HCR1 abundance declines during acclimation from VLC to HC.

To examine HCR1 localization, we imaged an HCR1-mGold-HA fusion expressed from the native *HCR1* promoter in the *hcr1* background. Under VLC, HCR1-mGold-HA-derived fluorescence was detected in the cytosol and along the two flagella; the flagellar signal was clearest in a focal plane optimized for the flagella (Fig. 1D). In the cell-body focal plane shown for HC, fluorescence was barely detectable when images were acquired and displayed with the same settings used for the corresponding VLC image. The VLC localization agrees with the flagellar-and-cytosolic localization reported for a *PsaD* promoter-driven LCI15-Venus fusion (35), whereas the native-promoter construct retains native CO_2_-responsive regulation.

### HCR1 represses the CCM under high CO_2_

To determine whether HCR1 contributes to CCM repression, we disrupted *HCR1* by inserting an *aphVII* hygromycin-resistance cassette at a Cas9-targeted site and confirmed the insertion by genotyping PCR (Fig. 2A and B). For complementation analysis, we introduced a wild-type genomic *HCR1* fragment into *hcr1*, generating a line hereafter designated *hcr1:HCR1*. We then used *hcr1* as the parental strain to disrupt *CBP1*, generating two independent *cbp1*/*hcr1* double-mutant lines. The *cbp1* single mutant was described previously (34).

**Fig. 2.**
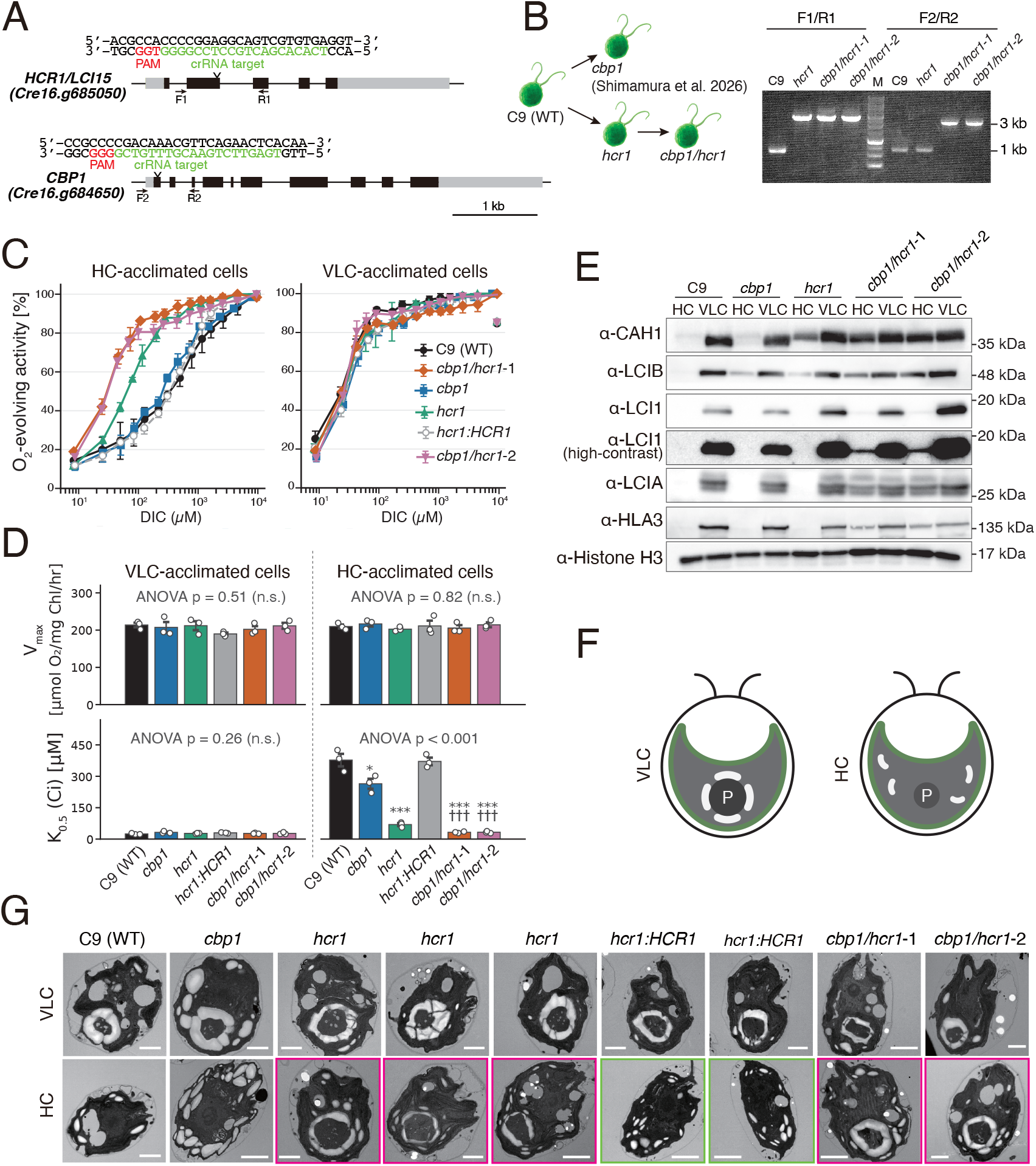
Generation and phenotypic characterization of *hcr1* and *cbp1*/*hcr1* mutants. **(A)** Gene models of *HCR1* (*Cre16.g685050*) and *CBP1* (*Cre16.g684650*), showing the crRNA targets, protospacer-adjacent motifs (PAMs), and genotyping primers. crRNA targets are green and PAMs are red. Black boxes, coding exons; gray boxes, untranslated regions; lines, introns. Scale bar, 1 kb. **(B)** Strain-construction scheme and genotyping PCR. F1/R1 and F2/R2 amplify the *HCR1* and *CBP1* loci, respectively. WT and marker-insertion alleles yielded products of approximately 1 and 3 kb, respectively. M, DNA size marker. **(C)** Light-saturated O_2_ evolution rates as a function of dissolved inorganic carbon (DIC) in HC- and VLC-acclimated cells. Rates were normalized to the value at 10 mM NaHCO_3_. Data are means ± SEM (*n* = 3 biological replicates). **(D)** V_max_ and K_0.5_(Ci) values derived from **(C)**. Bars are means ± SEM and circles are individual biological replicates. *P* values above the graphs are from one-way ANOVA. For HC K_0.5_(Ci), asterisks denote Holm-adjusted comparisons with C9 (\**P* < 0.05, \*\**P* < 0.01, \*\*\**P* < 0.001), and daggers denote comparisons with *hcr1* (†††*P* < 0.001). **(E)** Immunoblots of CAH1, LCIB, LCI1, LCIA, and HLA3 in HC- and VLC-acclimated cells. Histone H3 was the loading control. The high-contrast LCI1 panel is a uniformly adjusted rendering of the same blot shown immediately above. A representative result from three independent experiments is shown. **(F)** Schematic representation of pyrenoid and starch organization under VLC and HC. P, pyrenoid matrix. **(G)** Transmission electron micrographs of VLC- and HC-acclimated cells. Magenta frames mark the selected *hcr1* and *cbp1*/*hcr1* HC sections, and green frames mark the selected *hcr1*:*HCR1* HC sections. Scale bars, 2 µm.

To test whether loss of HCR1 derepresses the CCM under HC conditions, we measured light-saturated O_2_ evolution rates over a range of dissolved inorganic carbon (DIC) concentrations in HC- and VLC-acclimated cells and calculated V_max_ and K_0.5_(Ci), the DIC concentration required for half-maximal O_2_ evolution (Fig. 2C and D; Table S1). K_0.5_(Ci) values were log_10_-transformed for statistical testing. A two-way ANOVA revealed a significant genotype-by-acclimation-condition interaction for K_0.5_(Ci) (*F*_5,24_ = 113.4, *P* < 0.001). In HC-acclimated C9, K_0.5_(Ci) was 377.3 ± 29.9 µM. This value decreased to 264.2 ± 23.8 µM in *cbp1* (Holm-adjusted *P* = 0.018 versus C9) and to 69.4 ± 7.1 µM in *hcr1* (*P* < 0.001 versus C9), corresponding to 1.4- and 5.4-fold increases in apparent Ci affinity, respectively. The *hcr1:HCR1* complemented strain had a K_0.5_(Ci) of 371.0 ± 17.9 µM and did not differ from C9 (Holm-adjusted *P* = 0.91). The two *cbp1*/*hcr1* lines had still lower K_0.5_(Ci) values of 32.6 ± 2.0 and 32.9 ± 2.8 µM. Both values were significantly lower than that of *hcr1* (Holm-adjusted *P* = 1.2 × 10^−4^ for each comparison), representing a further 2.1-fold increase in apparent Ci affinity. Under VLC conditions, K_0.5_(Ci) ranged from 24.8 to 32.2 µM and did not differ among strains (one-way ANOVA on log₁₀-transformed values, P = 0.24). V_max_ did not differ among strains under either HC (*P* = 0.82) or VLC (*P* = 0.51) conditions. These results demonstrate that loss of *HCR1* specifically increases apparent Ci affinity under HC conditions and that additional loss of *CBP1* further strengthens this phenotype.

To determine whether the increased Ci affinity was accompanied by derepression of CCM proteins, we examined CAH1, LCIB, LCI1, LCIA, and HLA3 by immunoblotting (Fig. 2E). Under HC conditions, CAH1 and LCIB were undetectable or weak in C9, showed weaker accumulation in *cbp1*, and were readily detected in *hcr1* and both *cbp1*/*hcr1* lines. By contrast, LCI1, LCIA, and HLA3 were not detectably accumulated in *hcr1* under HC but showed clear signals in both double-mutant lines. Under VLC conditions, all five proteins were detected in all genotypes, although band intensities varied among strains. In the *hcr1* complemented line, CAH1 and LCIB returned to the C9-like pattern of weak or undetectable accumulation under HC and strong accumulation under VLC, whereas LCI1, LCIA, and HLA3 remained predominantly VLC associated (Fig. S1). Thus, loss of HCR1 alone was sufficient to derepress CAH1 and LCIB accumulation under HC, and this phenotype was reversed by HCR1 complementation. Detectable HC accumulation of the Ci-uptake proteins LCI1, LCIA, and HLA3 was associated with combined loss of *HCR1* and *CBP1*.

### HCR1 is required for high-CO_2_ remodeling of pyrenoid-associated starch

*Chlamydomonas reinhardtii* remodels pyrenoid-associated starch in response to CO_2_ availability. Under CO_2_-limiting conditions, curved starch plates accumulate around the Rubisco-rich pyrenoid matrix to form a sheath, whereas under HC conditions pyrenoid-associated starch is reduced and starch accumulates predominantly as granules in the chloroplast stroma (9, 36). These contrasting architectures are summarized schematically in Fig. 2F. To determine whether HCR1 affects this remodeling, we examined HC- and VLC-acclimated cells by transmission electron microscopy (Fig. 2G). Under VLC conditions, representative sections from all genotypes contained a conspicuous pyrenoid associated with curved starch plates. Under HC conditions, representative C9 and *cbp1* sections contained dispersed stromal starch granules and showed little or no organized pyrenoid-associated starch sheath. By contrast, three representative sections from *hcr1* and representative sections from both *cbp1*/*hcr1* lines retained a conspicuous pyrenoid partly or extensively surrounded by starch plates. This HC phenotype was not evident in the representative *hcr1:HCR1* section, consistent with complementation. Thus, the TEM images are consistent with incomplete HC-induced remodeling of pyrenoid-associated starch architecture in the absence of *HCR1*, paralleling the increased Ci affinity of *hcr1* and *cbp1*/*hcr1* cells.

### HCR1 selectively represses a core CCM transcriptional program under high CO_2_

To define the transcriptional effects of HCR1 and CBP1 under high CO_2_, we performed RNA-seq analysis. C9 was analyzed after acclimation to HC and VLC, whereas *cbp1*, *hcr1*, the *cbp1*/*hcr1*-1 double mutant, *cbp1:CBP1*, and *hcr1:HCR1* were analyzed under HC (Fig. 3A). Genes with CPM ≥ 1 in at least two libraries were retained, yielding 14,924 genes for analysis. Counts were normalized by the trimmed mean of M values (TMM), and differential expression was tested with the edgeR quasi-likelihood framework. Genes with FDR < 0.01 and |log_2_FC| ≥ 1 were classified as differentially expressed genes (DEGs). Mutant-versus-C9 comparisons were used to describe the total transcriptomic phenotype of each mutant, whereas direct comparisons between each mutant and its corresponding complemented line were used to reduce confounding by expression differences shared within a strain lineage (Fig. S2). We defined a CBP1- or HCR1-dependent DEG conservatively as a gene that met the DEG criteria and changed in the same direction in both the mutant-versus-C9 and mutant-versus-corresponding-complement comparisons.

**Fig. 3.**
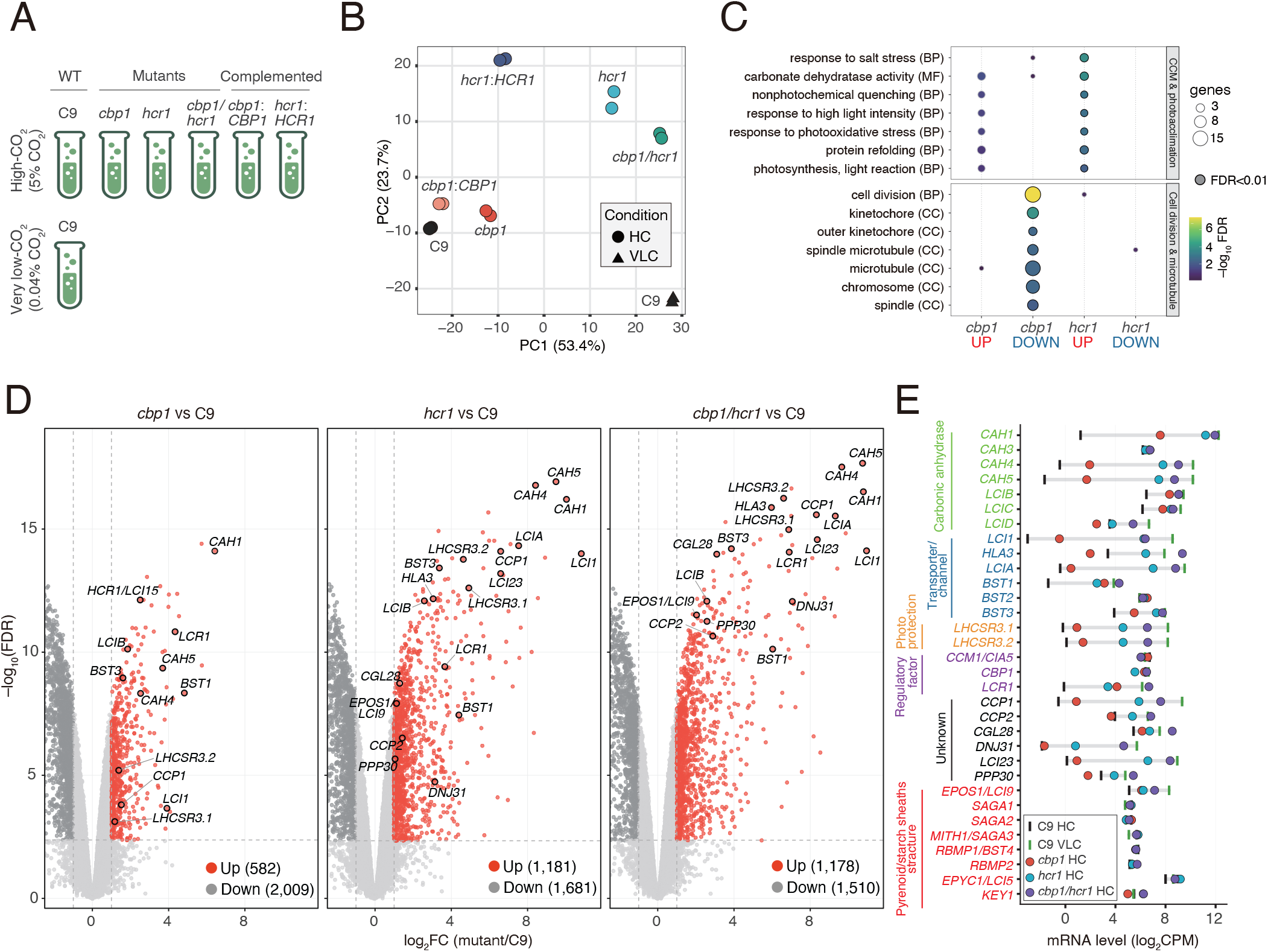
Transcriptome analysis of *cbp1*, *hcr1*, and *cbp1*/*hcr1* under HC. **(A)** RNA-seq design. C9 was analyzed under HC and VLC; *cbp1*, *hcr1*, *cbp1*/*hcr1*-1, *cbp1*:*CBP1*, and *hcr1*:*HCR1* were analyzed under HC (*n* = 2 biological replicates per genotype-condition group). **(B)** Principal component analysis of the C9 VLC-induced gene set. Points are biological replicates; colors denote genotypes and shapes denote HC (circles) or VLC (triangles). Text labels mark group centroids. PC1 and PC2 explain 53.4% and 23.7% of the variance, respectively. **(C)** GO over-representation analysis of complementation-validated DEGs. Columns denote the indicated up- or down-regulated gene sets. Dot size indicates the number of genes, fill indicates −log_10_(FDR), and black outlines indicate FDR < 0.01. BP, biological process; MF, molecular function; CC, cellular component. **(D)** Volcano plots for each mutant versus C9 under HC. Dashed lines indicate |log_2_FC| = 1 and FDR = 0.01. Up-regulated DEGs are red, down-regulated DEGs are dark gray, and other genes are light gray. Selected CCM-associated genes are outlined and labeled. The positive log_2_FC range is shown at an expanded scale. **(E)** Mean expression of selected genes from two biological replicates. Black and green ticks indicate C9 under HC and VLC, respectively, and gray segments connect these reference values. Colored circles indicate the HC-acclimated mutants as defined in the key. Gene-label colors denote functional classes.

Principal component analysis showed that PC1, which explained 53.4% of the variance, separated C9 cells acclimated to HC and VLC conditions. Under HC, the *cbp1*, *hcr1*, and *cbp1*/*hcr1* mutants were progressively displaced along PC1 toward VLC-acclimated C9, whereas the complemented lines shifted back toward HC-acclimated C9 (Fig. 3B). PC2, which explained 23.7% of the variance, separated *hcr1*, *hcr1:HCR1*, and *cbp1*/*hcr1* from the other strains. Because this separation persisted after HCR1 complementation, it likely represents a lineage-specific background component rather than an effect of HCR1 loss. We therefore defined complementation-validated DEGs as genes that changed in the same direction in both the mutant-versus-C9 and mutant-versus-complemented-line comparisons.

The complementation-validated sets comprised 204 up-regulated and 587 down-regulated CBP1-dependent genes and 66 up-regulated and 88 down-regulated HCR1-dependent genes. Gene Ontology (GO) analysis showed that HCR1-dependent up-regulated genes were enriched for carbonate dehydratase activity and photoacclimation-related terms, whereas CBP1-dependent down-regulated genes were enriched for cell-division, kinetochore, spindle, chromosome, and microtubule-related terms (Fig. 3C). No significant terms were identified for the CBP1-dependent up-regulated or HCR1-dependent down-regulated sets.

Volcano plots comparing each mutant with C9 under HC identified 582 up-regulated and 2,009 down-regulated DEGs in *cbp1*, 1,181 up-regulated and 1,681 down-regulated DEGs in *hcr1*, and 1,178 up-regulated and 1,510 down-regulated DEGs in *cbp1*/*hcr1* (Fig. 3D). Loss of HCR1 strongly increased the expression of core CCM and low-CO_2_-responsive genes, including *CAH1*, *CAH4*, *CAH5*, *HLA3*, *LCIA*, *LCI1*, *LCIB*, *BST1*, *BST3*, and *LHCSR3.1/3.2*. Comparison with the corresponding complemented lines reduced the total number of DEGs from 2,862 to 259 for *hcr1* and from 2,591 to 1,757 for *cbp1* (Fig. S1). Thus, the broad transcriptomic difference between *hcr1* and C9 contains a substantial lineage-specific component. Nevertheless, core CCM genes remained among the most strongly up-regulated genes in *hcr1* relative to *hcr1:HCR1*, demonstrating that their derepression was associated with HCR1 loss.

Absolute-expression analysis further showed that HCR1 loss preferentially affected a core subset of CCM genes (Fig. 3E). For example, *CAH1* expression increased from 1.2 log_2_ CPM in C9 under HC to 7.6 in *cbp1*, 11.2 in *hcr1*, and 11.9 in *cbp1*/*hcr1*, approaching the level in VLC-acclimated C9 cells (12.2 log_2_ CPM). Similar patterns were observed for *HLA3*, *LCIA*, *CAH4*, *CAH5*, and *CCP1*, whereas many other CCM-associated or pyrenoid-structural genes changed little. Notably, transcripts of *LCI1*, *LCIA*, and *HLA3* were elevated in *hcr1* under HC, although the corresponding proteins were not detectably accumulated in this mutant (Figs. 2E and 3E). Mean *LCI1* transcript abundance was similar in *hcr1* and *cbp1*/*hcr1*, whereas mean *LCIA* and *HLA3* transcript abundance was greater in the double mutant. Thus, transcriptional derepression of these Ci-uptake genes in *hcr1* did not by itself result in detectable accumulation of their protein products under HC. These results indicate that HCR1 selectively represses a core CCM transcriptional program, but that an additional regulatory step separates transporter-gene transcription from transporter-protein accumulation.

### HCR1 selectively controls CO_2_-dependent CAS relocation

CAS and LCIB exhibit distinct CO_2_-dependent localization patterns in *Chlamydomonas*. Under VLC, thylakoid-associated CAS becomes enriched within the pyrenoid along pyrenoid tubules, producing a compact spoke-like pattern, whereas LCIB forms a ring at the pyrenoid periphery; under HC, CAS is distributed over chloroplast thylakoids and LCIB is dispersed through the chloroplast stroma (Fig. 4A and C; 20–22, 30). To test whether these redistributions require HCR1 or CBP1, we examined both proteins by indirect immunofluorescence after exposing HC-acclimated cells for 2 h to VLC (0.04% CO_2_), HC (5% CO_2_), or, for CAS, VHC (15% CO_2_).

**Fig. 4.**
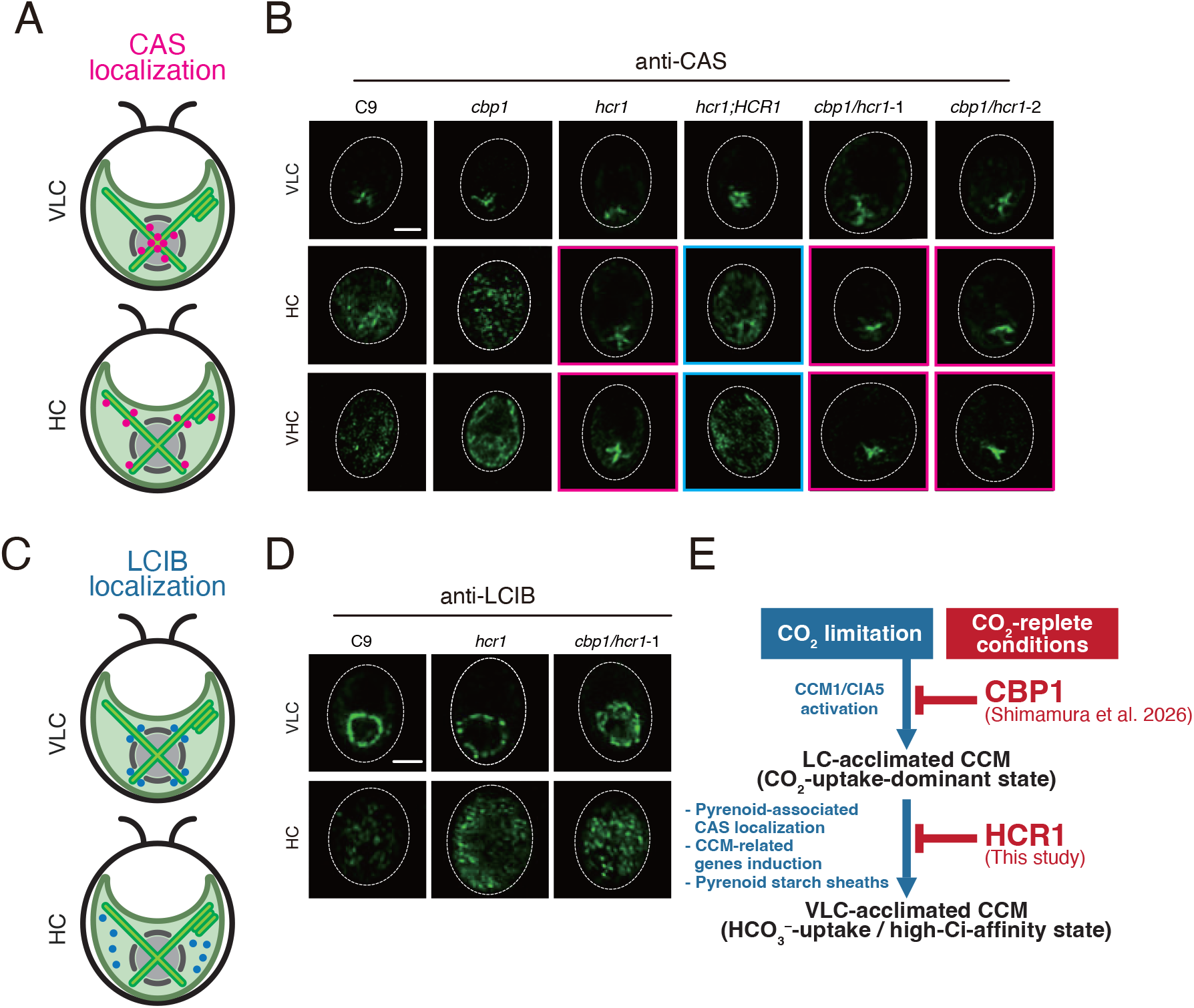
CO_2_-dependent localization of CAS and LCIB in *hcr1* and *cbp1*/*hcr1* mutants. **(A)** Schematic representation of CAS localization after 2 h under VLC and HC. CAS is magenta. Schematics are not to scale. **(B)** Indirect immunofluorescence of endogenous CAS in the indicated strains after 2 h under VLC (0.04% CO_2_), HC (5% CO_2_), or VHC (15% CO_2_). CAS is green and dashed lines indicate cell boundaries. Magenta frames mark persistent pyrenoid-associated CAS in HCR1-deficient cells under HC or VHC, and cyan frames mark restoration of the dispersed pattern in the complemented line. Scale bar, 5 µm. **(C)** Schematic representation of LCIB localization after 2 h under VLC and HC. LCIB is blue. Schematics are not to scale. **(D)** Indirect immunofluorescence of endogenous LCIB in the indicated strains after 2 h under VLC or HC. LCIB is green and dashed lines indicate cell boundaries. Scale bar, 5 µm. **(E)** Proposed model for the distinct roles of CBP1 and HCR1 in CO_2_-dependent CCM regulation. CO_2_ limitation activates CCM1/CIA5 to induce an LC-acclimated CCM, whereas CBP1 restrains this response under CO_2_-replete conditions. HCR1 is proposed to restrict the transition to a VLC-acclimated, high-Ci-affinity state associated with pyrenoid-localized CAS, starch-sheath development, and induction of the HCO_3_^−^ transporters HLA3 and LCIA. Blue arrows indicate induction or transition, and red bars indicate negative regulation.

In C9 and *cbp1*, CAS displayed the expected pyrenoid-associated pattern under VLC and a dispersed chloroplast pattern under HC and VHC (Fig. 4B). By contrast, *hcr1* and both independent *cbp1*/*hcr1* lines retained a compact pyrenoid-associated CAS signal under HC and VHC, as well as under VLC. The dispersed HC and VHC pattern was restored in the *hcr1:HCR1* complemented line. Thus, HCR1, but not CBP1 alone, is required for CO_2_-dependent redistribution of CAS away from the pyrenoid under CO_2_-replete conditions. Because the phenotype persisted after 2 h at 15% CO_2_, a threefold increase over the HC concentration, it is unlikely to reflect only a modest upward shift in the CO_2_ concentration required for CAS redistribution.

Persistent pyrenoid-associated CAS under HC paralleled the derepression of *HLA3* and *LCIA*, which were previously identified as CAS-dependent genes (Fig. 3). This association supports a model in which HCR1-dependent CAS redistribution contributes to suppression of CAS-associated chloroplast-to-nucleus signaling under HC, although localization alone does not establish that CAS causes the transcriptional phenotype.

LCIB showed a different result. In C9, *cbp1*, *hcr1*, and both *cbp1*/*hcr1* lines, LCIB formed a pyrenoid-peripheral ring under VLC and was dispersed throughout the chloroplast under HC (Fig. 4C and D). Thus, loss of HCR1 selectively disrupted the CO_2_-dependent localization of CAS without producing an obvious defect in LCIB redistribution. The preserved LCIB response argues against a general failure of CO_2_-responsive chloroplast protein relocation in *hcr1* and *cbp1*/*hcr1* cells.

## Discussion

### HCR1 defines a preloaded checkpoint for CCM shutdown

This study identifies HCR1 as a second negative regulator of the *Chlamydomonas* CCM with a function distinct from that of the nuclear repressor CBP1 (31). Under HC, loss of HCR1 produced a high-Ci-affinity phenotype, and additional loss of CBP1 shifted the double mutants further toward the VLC-acclimated state without changing maximum photosynthetic capacity (Fig. 2C and D). In *hcr1*, this physiological phenotype was accompanied by accumulation of CAH1 and LCIB and retention of pyrenoid-associated starch. The double mutants additionally accumulated LCI1, LCIA, and HLA3 under HC and showed a further increase in apparent Ci affinity (Fig. 2E–G). This stepwise progression indicates that HCR1 and CBP1 act as complementary checkpoints rather than redundant repressors.

Transcriptome analysis supports this distinction but also reveals a lineage-specific background component in the *hcr1*-derived strains. Comparison with the complemented line markedly reduced the broad differences observed against C9 (Fig. 3B and D; Fig. S1). The remaining HCR1-dependent genes were concentrated in CCM and photoacclimation functions, whereas CBP1-dependent changes extended to additional processes, including cell-division-related categories (Fig. 3C–E). HCR1 is therefore best viewed as a selective regulator of the high-affinity CCM rather than as a stronger global transcriptional repressor than CBP1.

The central mechanistic paradox is that HCR1 accumulates under VLC, whereas its mutant phenotype is expressed primarily under HC. HCR1 transcript and protein abundance were high under VLC, but HCR1-PA declined after transfer to HC and became barely detectable during prolonged HC acclimation (Fig. 1A and C). We propose that HCR1 is a preloaded reset factor rather than a stable component of the final HC state. CO_2_ repletion may activate the HCR1 pool accumulated during carbon limitation, allowing it to promote early redistribution of CAS away from pyrenoid tubules, terminate CAS-associated chloroplast-to-nucleus signaling, and facilitate remodeling of pyrenoid-associated starch. Rapid resetting would prevent continued expenditure on Ci uptake and the electron-transfer reactions that support lumen acidification when CO_2_ is abundant (23, 32, 33). Once the VLC program has been terminated, HCR1 may no longer be required and can decline. Thus, the inhibitory connection assigned to HCR1 in Fig. 4E denotes restriction of VLC-state persistence during CO_2_ repletion, not inhibition of VLC acclimation while CO_2_ remains limiting. Alternatively, a residual HCR1 pool may maintain repression, or transient HCR1 activity may establish a self-maintaining chloroplast state. Acute perturbation of HCR1 during controlled VLC-to-HC transitions will be required to distinguish these possibilities.

### HCR1 selectively connects CAS positioning to CCM repression

CAS showed the expected redistribution between pyrenoid tubules under VLC and thylakoid membranes throughout the chloroplast under HC in C9 and *cbp1* cells. In contrast, *hcr1* and *cbp1*/*hcr1* cells retained pyrenoid-associated CAS under HC and VHC, and complementation restored the dispersed pattern (Fig. 4A and B). LCIB nevertheless retained its normal CO_2_-responsive redistribution in the *hcr1* cells (Fig. 4C and D). HCR1 is therefore unlikely to be a general CO_2_ sensor or a universal regulator of chloroplast protein movement. It instead controls a CAS-specific branch or a structural state that selectively governs CAS positioning. The route from cytosolic HCR1 to thylakoid-associated CAS remains unknown. HCR1 could regulate a cytosolic chloroplast-signaling factor, a kinase or phosphatase, or a CAS-binding protein; alternatively, it may first remodel the pyrenoid, with CAS redistribution occurring secondarily. Its WW domain provides a plausible protein-interaction module (37), making identification of HCR1-interacting proteins a central mechanistic priority.

Genes previously reported to depend on CAS remained prominent among the complementation-validated genes derepressed in *hcr1*, including genes encoding Ci transporters, carbonic anhydrases, and photoacclimation proteins (Fig. 3D and E; Fig. S1). This correspondence is consistent with HCR1-dependent CAS redistribution contributing to termination of retrograde signaling, but it does not establish causality. Disruption of *CAS* in *hcr1* or *cbp1*/*hcr1* backgrounds will be required to test this model directly.

Transcript and protein accumulation were nevertheless uncoupled. Transcripts of *LCI1*, *LCIA*, and *HLA3* were strongly derepressed in *hcr1* under HC, but the corresponding proteins were not detectably accumulated until *CBP1* was also disrupted (Figs. 2E and 3E). For *LCIA* and particularly *HLA3*, the greater mean transcript abundance in the double mutant may contribute to a quantitative threshold for protein accumulation. This explanation is insufficient for *LCI1*, whose transcript abundance was similar in *hcr1* and *cbp1*/*hcr1*. The discrepancy therefore points to an additional regulatory layer involving translation, membrane insertion, intracellular trafficking, complex assembly, or protein stability. Because CAH1 and LCIB accumulated in *hcr1*, the discrepancy does not reflect a general failure of CCM-protein synthesis and may preferentially affect membrane-associated Ci-uptake proteins.

The present data do not show that CBP1 directly controls any of these post-transcriptional processes. A more conservative interpretation is that combined loss of *HCR1* and CBP1 completes the regulatory program required for stable accumulation of the Ci-uptake machinery. HCR1 loss may therefore prime the high-affinity CCM at the levels of gene transcription, CAS localization, and pyrenoid structure, whereas removal of both checkpoints permits robust accumulation of LCI1, LCIA, and HLA3. This distinction provides a plausible explanation for the further increase in apparent Ci affinity in the double mutants. Transcriptional derepression alone is therefore not a reliable proxy for assembly of a functional high-affinity CCM.

### HCR1 links high-affinity physiology to pyrenoid remodeling

HCR1 loss also altered pyrenoid-associated starch under HC. The *hcr1* and *cbp1*/*hcr1* sections retained conspicuous pyrenoid matrices associated with curved starch plates, whereas this architecture was not evident in the complemented line (Fig. 2F and G). A persistent starch sheath could enhance Ci affinity by restricting CO_2_ leakage and maintaining favorable geometry for localized CO_2_ delivery (9, 38), although quantitative morphometry is needed to establish penetrance. The causal order between CAS retention and starch persistence remains unresolved. Pyrenoid architecture can regulate CAS signaling, because disruption of *SAGA1* alters pyrenoid structure, CAS localization, and CAS-dependent gene expression (31, 39), but persistent CAS signaling could also stabilize a VLC-like pyrenoid state. The limited changes in known pyrenoid and starch-sheath genes argue against broad transcriptional induction of the structural program (Fig. 3E), suggesting post-transcriptional control, altered starch metabolism, or remodeling of protein interactions. Moreover, retained starch plates did not recruit LCIB under HC (Figs. 2G and 4D), even though pyrenoid-associated starch is required for normal LCIB recruitment under VLC (9). CAS positioning, starch organization, and LCIB relocation are therefore separable modules rather than outputs of a single global switch.

### CBP1 and HCR1 form complementary checkpoints

CBP1 and HCR1 share CobW_C and WW domains but differ in domain composition, localization, and output. Nuclear CBP1 interacts with CCM1 and restrains CCM1-dependent transcription (34), whereas cytosolic HCR1 preferentially controls CAS positioning, pyrenoid remodeling, and the high-affinity CCM. The enrichment of cell-division-related terms among CBP1-dependent down-regulated genes may reflect the energetic burden of inappropriate CCM activity rather than direct CBP1 control of the cell cycle (23, 32, 33). Their related architectures suggest functional divergence from a shared regulatory module: CobW-family proteins are metal-handling P-loop GTPases, whereas WW domains mediate protein interactions (37, 40). Whether these domains explain the nuclear and cytosolic specialization of CBP1 and HCR1 remains speculative.

The model in Fig. 4E integrates these findings. CO_2_ limitation activates CCM1 and establishes an LC-acclimated, CO_2_-uptake-dominant program (24–27). More severe limitation engages pyrenoid-associated CAS, starch-sheath development, HLA3/LCIA-dependent HCO_3_^−^ uptake, and the high-affinity VLC physiology (9, 10, 12, 20, 21, 23, 30, 31). Under CO_2_-replete conditions, CBP1 restrains CCM1-dependent transcription at a nuclear checkpoint, whereas HCR1 is proposed to terminate persistence of the VLC-specific branch by enabling CAS redistribution and pyrenoid-starch remodeling. The double-mutant phenotype further suggests that full engagement of this branch requires both transcriptional derepression and a separate step that permits stable accumulation of the Ci-uptake machinery. This two-checkpoint model explains the partial phenotype of *cbp1*, the stronger high-affinity and structural phenotypes of *hcr1*, and the near-VLC state of the double mutant under HC (Figs. 2–4). It also reframes CCM shutdown as an active resetting process. The key unresolved questions are what activates HCR1 upon CO_2_ repletion, how cytosolic HCR1 communicates with CAS, and whether CAS relocation causes or follows pyrenoid remodeling.

## Materials and Methods

### *Chlamydomonas reinhardtii* strains and culture conditions

The wild-type strain was C9 (CC-5098, *mt^−^*). The *cbp1*-1 mutant, hereafter *cbp1*, and the *cbp1:CBP1* complemented line were described previously (34). Additional strains were *hcr1*, two independent *cbp1*/*hcr1* double mutants, and the complemented or tagged lines *hcr1:HCR1*, *hcr1:HCR1-PA*, and *hcr1:HCR1-mGold-PA*. Cells were maintained in Tris-acetate-phosphate medium at 25°C under continuous white light and transferred to MOPS-buffered phosphate medium for experiments. Cultures were aerated with 5% CO_2_ in air for high-CO_2_ conditions, ambient air containing approximately 0.04% CO_2_ for very-low-CO_2_ conditions, or 15% CO_2_ in air for very-high-CO_2_ conditions. These conditions are referred to as HC, VLC, and VHC, respectively. Air-aerated liquid cultures were designated VLC because photosynthetic CO_2_ consumption lowers dissolved CO_2_ below approximately 7 µM in this system (20, 23).

### Phylogenetic and domain analyses

The 14 COG0523/CobW-related proteins encoded by *C. reinhardtii* were identified from the Phytozome v5.5 proteome by HMMER searches against the Pfam CobW and CobW_C profiles. Full-length sequences were aligned using MAFFT version 7, and a neighbor-joining tree based on uncorrected amino acid p-distances was inferred in MEGA X with 1,000 bootstrap replicates and was midpoint-rooted. CobW, CobW_C, and WW domains were annotated using the corresponding Pfam profiles.

### Generation of mutant, complemented, and tagged strains

The *hcr1* mutant was generated by Cas9-crRNA ribonucleoprotein-mediated insertion of the *aphVII* cassette into C9. Two independent *cbp1*/*hcr1* lines were generated by disrupting *CBP1* in the *hcr1* background with an *aphVIII* cassette and were verified by locus-specific PCR and Sanger sequencing. Native-promoter genomic constructs were introduced by electroporation to generate *hcr1:HCR1*, *hcr1:HCR1-PA*, and *hcr1:HCR1-mGold-PA*. The untagged complemented line was verified by genotyping and recovery of CCM suppression under HC conditions.

### Confocal imaging and immunoblot analysis

HCR1-mGold-PA localization was examined with a Leica TCS SP8 confocal laser-scanning microscope. mGold was excited at 514 nm and detected at 520 to 550 nm, and chlorophyll autofluorescence was detected at 648 to 700 nm. Condition-matched images were acquired with identical settings and deconvolved using Huygens Essential. For immunoblotting, protein loading was normalized to chlorophyll content, proteins were separated by SDS-PAGE and transferred to polyvinylidene difluoride membranes, and HCR1-PA or CCM proteins were detected with the antibodies specified in SI Appendix.

### Photosynthetic O_2_ evolution and statistical analysis

Light-saturated O₂ evolution was measured with a Clark-type electrode in Ci-depleted HEPES buffer while NaHCO_3_ was added stepwise. The maximum O_2_ evolution rate, V_max_, and the DIC concentration required for half-maximal O_2_ evolution, K_0.5_(Ci), were obtained from the Ci-response curves. Three independently cultured biological replicates were analyzed for each genotype and condition. V_max_ was analyzed on the original scale, whereas K_0.5_(Ci) was log_10_-transformed. Genotype and acclimation-condition effects were evaluated by two-way ANOVA, followed by condition-specific one-way ANOVA and Holm-adjusted planned contrasts where appropriate.

### Transmission electron microscopy

Cells were fixed with paraformaldehyde and glutaraldehyde, postfixed with osmium tetroxide, dehydrated, embedded in epoxy resin, and sectioned for transmission electron microscopy with modifications to the method as described previously (9). Ultrathin sections were stained with uranyl acetate and lead citrate and examined using a JEM-1400Flash transmission electron microscope.

### RNA-seq and bioinformatic analyses

Total RNA was purified with the RNeasy Plant Mini Kit, and libraries from two independent biological replicates per genotype-condition group were sequenced on an Illumina NextSeq 500 using a High Output 75-cycle kit. Reads were trimmed with Trim Galore!, aligned to the *C. reinhardtii* v5.6 genome with HISAT2, and counted against the Phytozome v5.5 annotation with htseq-count. Differential expression was analyzed in edgeR after TMM normalization. Genes with CPM ≥ 1 in at least two libraries were retained, and genes with FDR < 0.01 and |log_2_FC| ≥ 1 were classified as differentially expressed. Complementation-validated genes changed in the same direction in both mutant-versus-C9 and mutant-versus-corresponding-complement comparisons. Principal component analysis used the C9 VLC-induced gene set, and GO enrichment was evaluated by one-sided hypergeometric tests with Benjamini-Hochberg correction.

### Indirect immunofluorescence microscopy

Indirect immunofluorescence was performed as described previously (41). Endogenous CAS or LCIB was detected with rabbit primary antibodies and an Alexa Fluor 488-conjugated secondary antibody. Images were acquired with a Leica TCS SP8 using identical settings within each antibody series and were processed identically.

Detailed procedures for all experiments described above are provided in SI Appendix.

### Data Availability

The raw RNA-seq data generated in this study have been deposited and are publicly available in the DDBJ Sequence Read Archive under BioProject accession PRJDB43043 (https://ddbj.nig.ac.jp/search/entry/bioproject/PRJDB43043) (42), with Run accessions DRR1090280–DRR1090293. The associated sample metadata are available under BioSample accessions SAMD01960727–SAMD01960740. Strains generated in this study will be available from the Chlamydomonas Resource Center (https://www.chlamycollection.org/) (43) upon publication. The accession numbers of the Phytozome database for Chlamydomonas gene *HCR1* is *Cre16.g685050* (44). All other data are included in the manuscript and/or SI Appendix.

## Acknowledgments

We thank Hiroko Shimada, Kazuko Tarui, Nahoko Nishimura, and Hatsue Mizuhara for technical assistance. We thank Yoshihiro Yoshitake and Yukari Sando at the Next-Generation Sequencing Facility, Graduate School of Biostudies, Kyoto University, for support with RNA-seq. We thank Keiko Okamoto-Furuta and Haruyasu Kohda at the Division of Electron Microscopic Study, Center for Anatomical Studies, Graduate School of Medicine, Kyoto University, for assistance with electron microscopy.

## SI Appendix

### Supplementary Materials and Methods

#### *Chlamydomonas reinhardtii* strains and culture conditions

The wild-type *Chlamydomonas reinhardtii* strain used in this study was C9 (CC-5098, *mt^−^*), originally obtained from the IAM Culture Collection, University of Tokyo, and currently available as NIES-2235 and CC-5098. The *cbp1*-1 mutant, hereafter referred to as *cbp1*, and the *cbp1:CBP1* complemented line were described previously (1). Additional strains analyzed in this study were the *hcr1* mutant, two independently isolated *cbp1*/*hcr1* double-mutant lines designated *cbp1*/*hcr1*-1 and *cbp1*/*hcr1*-2, and the complemented or tagged lines *hcr1:HCR1*, *hcr1:HCR1-PA*, and *hcr1:HCR1-mGold-PA*. The generation and validation of these strains are described below.

Cells were pre-cultured in 5 mL Tris-acetate-phosphate (TAP) medium for at least 12 h at 25°C under continuous white light at approximately 120 µmol photons m^−2^ s^−1^ with orbital agitation. For photoautotrophic experiments, cells were transferred to 50 mL MOPS-buffered phosphate (MOPS-P) medium, consisting of phosphate medium supplemented with 20 mM MOPS at pH 7.0. Cultures were continuously aerated with 5% CO_2_ in air for high-CO_2_ conditions, ambient air containing approximately 0.04% CO_2_ for very-low-CO_2_ conditions, or 15% CO_2_ in air for very-high-CO_2_ conditions. These conditions are hereafter referred to as HC, VLC, and VHC, respectively. Experimental cultures were maintained in mid-logarithmic growth at an optical density at 730 nm of approximately 0.3 to 0.6 to minimize self-shading and gas limitation.

Ambient-air aeration of liquid cultures was designated as VLC because photosynthetic CO_2_ consumption lowers the dissolved CO_2_ concentration below approximately 7 µM after acclimation, as indicated by the pyrenoid-peripheral localization of LCIB (2, 3). This classification is based on previous measurements in the same liquid-culture system, in which dissolved inorganic carbon (Ci) was quantified by gas chromatography after methanization and the dissolved CO_2_ concentration was calculated using the Henderson– Hasselbalch equation.

Unless otherwise stated, cells were first acclimated to HC for at least 12 h, collected by centrifugation at 600 × g for 5 min, resuspended in fresh MOPS-P medium, and transferred to the indicated CO_2_ condition. For the HCR1-PA abundance time course, VLC-acclimated *hcr1:HCR1-PA* cells were transferred to HC and sampled at 0, 0.5, 1, 2, 4, 8, and 12 h. For CAS immunofluorescence, HC-acclimated cells were transferred for 2 h to VLC, HC, or VHC. For LCIB immunofluorescence, HC-acclimated cells were transferred for 2 h to VLC or HC.

#### Identification of COG0523/CobW-related proteins, phylogenetic analysis, and domain annotation

The 14-protein COG0523/CobW-related set was defined as all proteins in the predicted proteome of *C. reinhardtii* (Phytozome, Creinhardtii v5.5) that scored above the family-specific gathering threshold for either the CobW or CobW_C Pfam profile. Full-length amino-acid sequences were aligned using MAFFT version 7 (4). A neighbor-joining tree was inferred in MEGA X (5) from uncorrected pairwise amino-acid p-distances, with pairwise deletion of alignment gaps and 1,000 bootstrap replicates and was midpoint-rooted for display. The transcript-abundance columns in Fig. 1A show mean TMM-normalized log_2_(CPM + 1) values from the two C9 HC and two C9 VLC RNA-seq libraries described below.

CobW, CobW_C, and WW domains were annotated using the Pfam profile hidden Markov models PF02492, PF07683, and PF00397 obtained from InterPro (https://www.ebi.ac.uk/interpro/). Protein sequences were searched using HMMER 3.4 through pyhmmer v0.12.1 with family-specific gathering thresholds. Adjacent hits to the same profile separated by ≤ 50 aa were merged into a single domain box for display. Gene coordinates and exon-intron structures were obtained from the Phytozome v5.5 annotation.

#### Generation and genotyping of mutant strains

The *cbp1*-1 mutant was generated previously by Cas9-crRNA ribonucleoprotein (RNP)-mediated insertion of the hygromycin-resistance cassette *aphVII* into the second exon of *CBP1* (1). The *hcr1* mutant was generated in C9 by Cas9 RNP-mediated insertion of *aphVII* at the target shown in Fig. 2A. Disruption of *HCR1* was verified by locus-specific PCR.

To generate double mutants, a crRNA targeting the first exon of *CBP1* was designed with CRISPOR (6, 7). Candidates with a Doench score ≥ 60 and few predicted off-target sites were prioritized. Synthetic crRNA and tracrRNA (Integrated DNA Technologies) were assembled with Cas9 to form an RNP and co-electroporated into *hcr1* cells with a PCR-amplified *aphVIII* cassette conferring paromomycin resistance, using the electroporation procedure described previously (8). Cells were recovered for 12–18 h at 25°C under approximately 1.5 µmol photons m^−2^ s^−1^ and plated on MOPS-P agar containing 30 µg mL⁻¹ paromomycin. Plates were incubated for 4 d at 25°C under approximately 45 µmol photons m^−2^ s^−1^. Two independently isolated double-mutant lines were retained.

For colony PCR, cells from individual colonies were suspended in 50 µL of 5% Chelex 100 resin, vortexed for 10 s, heated at 100°C for 10 min, cooled on ice for 1 min, and centrifuged at 15,000 × g for 1 min at 4°C. The supernatant was used as the PCR template. In *cbp1*/*hcr1*-1 and *cbp1*/*hcr1*-2, a single *aphVIII* cassette insertion at the CBP1 target site was confirmed by Sanger sequencing.

#### Generation of complemented and tagged HCR1 lines

To generate the untagged complementation line *hcr1:HCR1*, a genomic fragment encompassing 1,229 bp upstream of the *HCR1* translational start site, the complete coding region, and 1,389 bp downstream of the stop codon was introduced into *hcr1* cells by electroporation. Transformants were selected for paromomycin resistance conferred by the *aphVIII* marker. Complementation was confirmed by genotyping and restoration of CCM suppression under HC conditions.

To generate *hcr1:HCR1-PA* for the HCR1 abundance time course, HCR1 was fused in frame at its C terminus to a PA epitope tag and expressed under the control of its native promoter in the *hcr1* background. To generate *hcr1:HCR1-mGold-PA* for subcellular localization analysis, a construct comprising approximately 2 kb of the native *HCR1* upstream region, the complete *HCR1* coding sequence, and an in-frame C-terminal mGold-PA fusion was introduced into *hcr1* cells by electroporation.

#### Confocal imaging of HCR1-mGold-PA

A 5-µL aliquot of cell suspension was placed on a coverslip and immobilized beneath a 1.5% agarose pad prepared in TAE buffer. Fluorescence was recorded with a Leica TCS SP8 confocal laser-scanning microscope. mGold was excited at 514 nm and detected at 520–550 nm; chlorophyll autofluorescence was detected at 648–700 nm. Images were acquired with 1.0% laser power and a detector gain of 300%. Condition-matched cell-body images were acquired with identical settings. Raw image stacks were deconvolved using Huygens Essential (Scientific Volume Imaging). Scale-bar calibration was obtained from the microscope metadata.

#### Immunoblot analysis

Cells were harvested, and total proteins were extracted, denatured, separated by SDS-PAGE, and transferred to polyvinylidene difluoride membranes as described previously (1, 9). Sample loading was normalized to chlorophyll content. For analysis of HLA3, LCIA, and LCI1, extracts corresponding to 2 µg chlorophyll were loaded per lane. For analysis of CAH1 and LCIB, extracts corresponding to 1 µg chlorophyll were loaded per lane. Extracts corresponding to 1 µg chlorophyll were also loaded for the HCR1-PA time-course experiment.

The primary antibodies used were rabbit anti-LCIB at 1:5,000 (10), rabbit anti-CAH1/2 at 1:2,500 (11), rabbit anti-LCIA at 1:5,000 (12), rabbit anti-HLA3 at 1:1,250 (12), rabbit anti-LCI1 at 1:5,000 (13), rabbit anti-histone H3 at 1:20,000 (Abcam), rabbit anti-CBP1 at 1:10,000 (1), and rat anti-PA at 1:5,000 (FUJIFILM). Horseradish peroxidase-conjugated goat anti-rabbit IgG at 1:10,000 (Life Technologies) or goat anti-rat IgG at 1:2,000 (R&D Systems) was used as the secondary antibody, as appropriate. Chemiluminescent signals were developed using Luminata Crescendo Western HRP substrate and captured with an ImageQuant LAS 4010 imaging system.

Any linear adjustment of image brightness or contrast was applied uniformly to the entire image. The high-contrast LCI1 panel in Fig. 2E was generated from the same source image as the standard-contrast panel and was included solely to visualize weak LCI1 signals, not for quantitative comparison.

#### Measurement of photosynthetic O₂ evolution and Ci-response kinetics

HC- or VLC-acclimated cells were collected at 600 × g for 5 min and resuspended in 50 mM HEPES (pH 7.8) to a chlorophyll concentration of 10–20 µg mL⁻¹. Cell suspensions were depleted of inorganic carbon by bubbling with N_2_. O₂ evolution was measured with a Clark-type oxygen electrode (Hansatech Instruments) under 750 µmol photons m^−2^ s^−1^ white light. NaHCO_3_ was added stepwise at 30-s intervals, and the O_2_ signal was recorded with an LR4220 recorder (Yokogawa). K_0.5_(Ci) is the DIC concentration yielding half-maximal O_2_ evolution. For Fig. 2C, each rate was normalized to the rate measured at 10 mM NaHCO_3_; absolute fitted V_max_ and K_0.5_(Ci) values are shown in Fig. 2D. Each biological replicate was an independently cultured and acclimated sample.

#### Transmission electron microscopy

Cells were prepared for transmission electron microscopy with modifications to the method as described previously (14). Cultures were harvested by centrifugation at 600 × g for 5 min at room temperature. Cell pellets were resuspended in 0.1 M phosphate buffer at pH 7.4 containing 4% paraformaldehyde and 2% glutaraldehyde and fixed on ice for 4 h, with gentle mixing once per hour. After washing with 0.1 M phosphate buffer, the fixed cells were embedded in SeaPlaque low-melting-point agarose and incubated overnight at 4°C in the same fixative.

Subsequent sample processing was performed by the Electron Microscopy Facility of the Graduate School of Biostudies, Kyoto University. Agarose-embedded samples were postfixed with 1% osmium tetroxide in 0.1 M phosphate buffer for 3 h at room temperature and dehydrated through a graded ethanol series. Samples were transferred to propylene oxide and sequentially infiltrated with mixtures of propylene oxide and LUVEAK-812 at ratios of 3:1, 1:1, and 1:3 for 90 min each. The samples were then embedded in Epon 812 resin and polymerized. Ultrathin sections were stained with uranyl acetate and lead citrate and examined using a JEM-1400Flash transmission electron microscope (JEOL).

#### RNA extraction, library preparation, and sequencing

Cells were harvested by centrifugation at 600 × g for 5 min at room temperature and resuspended in 200 µL TE buffer. The cell suspension was frozen dropwise in liquid N_2_ and disrupted using a prechilled mortar and pestle. Total RNA was treated with DNase I and purified using the RNeasy Plant Mini Kit (QIAGEN).

RNA concentration was measured using a Qubit fluorometer, and RNA quality was assessed using a Bioanalyzer at the Next-Generation Sequencing Facility, Graduate School of Biostudies, Kyoto University. RNA-seq libraries were prepared by the facility. Following library preparation, library concentration and fragment-size distribution were evaluated using Qubit fluorometry and a Bioanalyzer, and library concentrations were further determined by qPCR. Sequencing was performed in a single run on an Illumina NextSeq 500 using a High Output 75-cycle kit. Two independently cultured biological replicates were sequenced for each genotype-condition group shown in Fig. 3A.

#### Read processing, gene-level quantification, and differential-expression analysis

Adapter sequences and low-quality bases were removed from the raw reads using Trim Galore!. The trimmed reads were aligned to the *Chlamydomonas reinhardtii* reference genome assembly v5.6 using HISAT2 (15). Gene-level read counts were generated using htseq-count in HTSeq (16), with gene models from the Phytozome v5.5 annotation. Gene identifiers therefore retain the “.v5.5” annotation suffix. The resulting count matrix contained 17,741 annotated gene models across 14 libraries, comprising two independently cultured biological replicates for each genotype and CO₂ condition. After exclusion of the htseq-count summary categories, the number of reads assigned to annotated gene models ranged from 18.3 million to 26.6 million per library, with a median of 21.6 million.

Differential expression was analyzed using edgeR 4.0.16 (17) under R 4.3.3. Genes with a counts-per-million value of at least 1 in at least 2 of the 14 libraries were retained, leaving 14,924 genes. Library sizes were normalized by the trimmed mean of M values (TMM). Dispersion estimation and quasi-likelihood fitting were performed with robust estimation (estimateDisp followed by glmQLFit with robust = TRUE). The design matrix contained one coefficient for each genotype-condition group (seven groups, fitted without an intercept), and all comparisons were extracted as contrasts from this single model. The analyzed contrasts were C9 VLC versus C9 HC, *cbp1* versus C9 under HC, *hcr1* versus C9 under HC, *cbp1*/*hcr1*-1 versus C9 under HC, *cbp1* versus *cbp1:CBP1* under HC, and *hcr1* versus *hcr1:HCR1* under HC. P values were adjusted using the Benjamini–Hochberg method. Genes with FDR < 0.01 and |log_2_FC| ≥ 1 were classified as differentially expressed genes.

A complementation-validated CBP1- or HCR1-dependent DEG was defined as a gene that met the DEG criteria and changed in the same direction in both the mutant-versus-C9 and mutant-versus-corresponding-complement comparisons. Mutant-versus-C9 contrasts were retained to describe the complete mutant transcriptomic phenotype, whereas the complementation criterion was used to reduce confounding by lineage-specific expression differences.

For Fig. 3E, TMM-normalized log_2_ CPM values were averaged across the two biological replicates of each genotype–condition group without further scaling, and the group means for *cbp1*, *hcr1*, and *cbp1*/*hcr1*-1 under HC were plotted against the C9 HC and C9 VLC means for the indicated genes.

#### Principal component and GO enrichment analyses

For Fig. 3B, the input set comprised 671 genes up-regulated in C9 under VLC relative to HC at FDR < 0.01 and log_2_FC ≥ 1. PCA was performed on TMM-normalized log_2_ CPM values (computed with cpm(log = TRUE, prior.count = 2)) using prcomp, with genes mean-centered and not variance-scaled; PC1 and PC2 accounted for 53.4% and 23.7% of the total variance. Replicate-level scores were plotted, and text labels indicate group centroids.

GO over-representation was tested separately for the complementation-validated CBP1-up, CBP1-down, HCR1-up, and HCR1-down gene sets using a one-sided hypergeometric test. The 14,924 genes retained for edgeR analysis were used as the background. Only terms annotated to at least three background genes and overlapping the query set by at least one gene were tested. GO annotations were downloaded from Ensembl Plants BioMart. FDR values were calculated using Benjamini–Hochberg correction across the tested terms separately for each of the four gene sets, with the three ontologies (BP, MF, and CC) corrected together within each set.

#### Indirect immunofluorescence microscopy

Indirect immunofluorescence was performed as described previously (18). After 2 h under the indicated CO_2_ condition, cells were fixed and permeabilized in parallel. Endogenous CAS or LCIB was detected with rabbit anti-CAS at 1:500 (9) or rabbit anti-LCIB primary antibody at 1:500 (12), followed by Alexa Fluor 488-conjugated goat anti-rabbit IgG (Life Technologies; 1:500). Fluorescence was acquired with a Leica TCS SP8 using 488-nm excitation, 520–550-nm detection, 1.0% laser power, and detector gain of 300%. Images were deconvolved using Huygens Essential. All samples within an antibody series were acquired using identical settings and processed identically. Paired differential-interference-contrast images were used to identify cell boundaries and the pyrenoid region.

## Supplementary Figures Legends

**Fig. S1.**
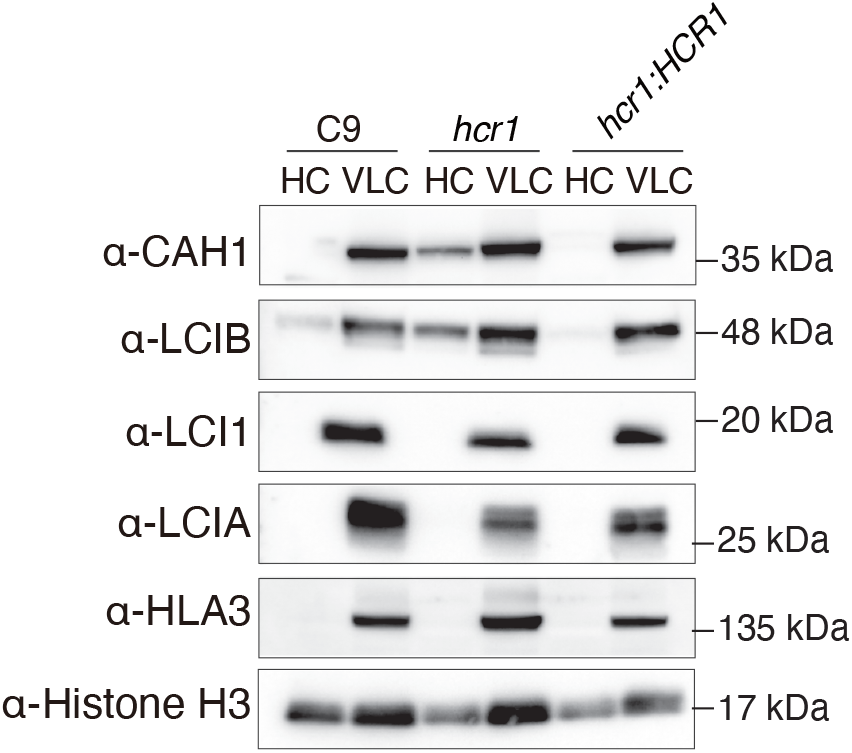
CCM-protein accumulation in C9, *hcr1*, and the *HCR1*-complemented line. Immunoblots of CAH1, LCIB, LCI1, LCIA, and HLA3 in C9, *hcr1*, and *hcr1:HCR1* cells acclimated to HC or VLC conditions. Histone H3 served as the loading control. Molecular-mass marker positions are indicated on the right.

**Fig. S2.**
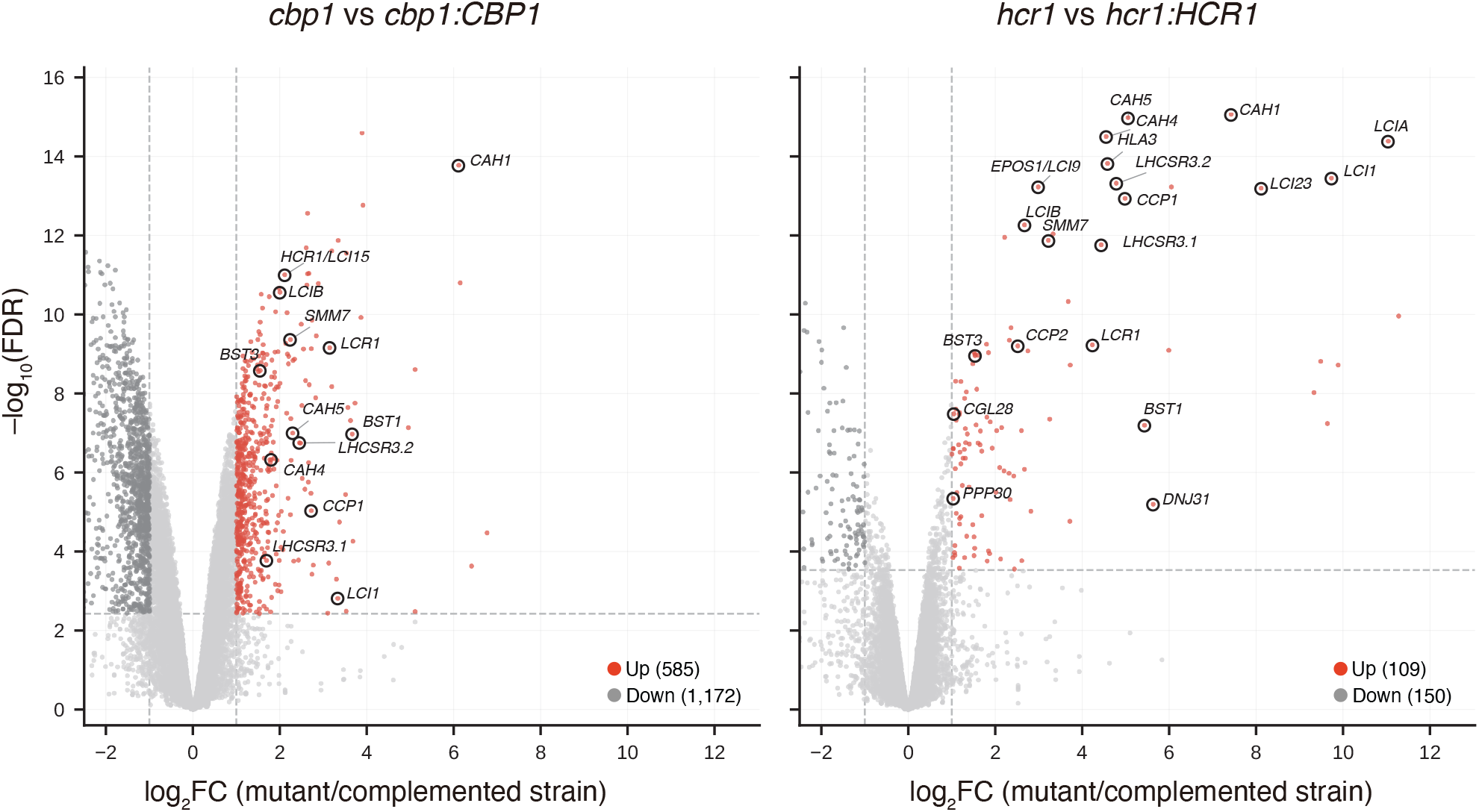
Volcano plots for each mutant versus complemented strain under HC. Dashed lines indicate |log₂FC| = 1 and FDR = 0.01. Up-regulated DEGs are red, down-regulated DEGs are dark gray, and other genes are light gray. Selected CCM-associated genes are outlined and labeled. The positive log2FC range is expanded.

## References

1. Giordano, M., Beardall, J. & Raven, J. A. CO_2_ concentrating mechanisms in algae: mechanisms, environmental modulation, and evolution. Annu Rev Plant Biol 56, 99–131 (2005).

2. Reinfelder, J. R. Carbon concentrating mechanisms in eukaryotic marine phytoplankton. Annu Rev Mar Sci 3, 291–315 (2011).

3. Raven, J. A., Beardall, J. & Giordano, M. Energy costs of carbon dioxide concentrating mechanisms in aquatic organisms. Photosynth Res 121, 111–124 (2014).

4. Barrett, J., Girr, P. & Mackinder, L. C. M. Pyrenoids: CO_2_-fixing phase separated liquid organelles. Biochim Biophys Acta Mol Cell Res 1868, 118949 (2021).

5. He, S., Crans, V. L. & Jonikas, M. C. The pyrenoid: the eukaryotic CO_2_-concentrating organelle. Plant Cell 35, 3236–3259 (2023).

6. Mackinder, L. C. M. et al. A repeat protein links Rubisco to form the eukaryotic carbon-concentrating organelle. Proc Natl Acad Sci USA 113, 5958–5963 (2016).

7. Freeman Rosenzweig, E. S., et al. The eukaryotic CO_2_-concentrating organelle is liquid- like and exhibits dynamic reorganization. Cell 171, 148–162.e19 (2017).

8. He, S. et al. The structural basis of Rubisco phase separation in the pyrenoid. Nat Plants 6, 1480–1490 (2020).

9. Toyokawa, C., Yamano, T. & Fukuzawa, H. Pyrenoid starch sheath is required for LCIB localization and the CO_2_-concentrating mechanism in green algae. Plant Physiol 182, 1883–1893 (2020).

10. Duanmu, D., Miller, A. R., Horken, K. M., Weeks, D. P. & Spalding, M. H. Knockdown of limiting-CO_2_-induced gene HLA3 decreases HCO_3_^−^ transport and photosynthetic Ci affinity in Chlamydomonas reinhardtii. Proc Natl Acad Sci USA 106, 5990–5995 (2009).

11. Gao, H., Wang, Y., Fei, X., Wright, D. A. & Spalding, M. H. Expression activation and functional analysis of HLA3, a putative inorganic carbon transporter in Chlamydomonas reinhardtii. Plant J 82, 1–11 (2015).

12. Yamano, T. et al. Characterization of cooperative bicarbonate uptake into chloroplast stroma in the green alga Chlamydomonas reinhardtii. Proc Natl Acad Sci USA 112, 7315–7320 (2015).

13. Förster, B. et al. The Chlamydomonas reinhardtii chloroplast envelope protein LCIA transports bicarbonate in planta. J Exp Bot 74, 3651–3666 (2023).

14. Guo, J. et al. Structure of Chlamydomonas reinhardtii LciA guided the engineering of FNT family proteins to gain bicarbonate transport activity. Nat Plants 12, 231–240 (2026).

15. Karlsson, J. et al. A novel alpha-type carbonic anhydrase associated with the thylakoid membrane in Chlamydomonas reinhardtii is required for growth at ambient CO_2_. EMBO J 17, 1208–1216 (1998).

16. Mukherjee, A. et al. Thylakoid localized bestrophin-like proteins are essential for the CO_2_ concentrating mechanism of Chlamydomonas reinhardtii. Proc Natl Acad Sci USA 116, 16915–16920 (2019).

17. Kono, A. & Spalding, M. H. LCI1, a Chlamydomonas reinhardtii plasma membrane protein, functions in active CO_2_ uptake under low CO_2_. Plant J 102, 1127–1141 (2020).

18. Shimamura, D. et al. Periplasmic carbonic anhydrase CAH1 contributes to high inorganic carbon affinity in Chlamydomonas reinhardtii. Plant Physiol 196, 2395–2404 (2024).

19. Wang, Y. & Spalding, M. H. Acclimation to very low CO_2_: contribution of limiting-CO_2_ inducible proteins, LCIB and LCIA, to inorganic carbon uptake in Chlamydomonas reinhardtii. Plant Physiol 166, 2040–2050 (2014).

20. Yamano, T., Toyokawa, C., Shimamura, D., Matsuoka, T. & Fukuzawa, H. CO_2_-dependent migration and relocation of LCIB, a pyrenoid-peripheral protein in Chlamydomonas reinhardtii. Plant Physiol 188, 1081–1094 (2022).

21. Yamano, T. et al. Light and low-CO_2_-dependent LCIB–LCIC complex localization in the chloroplast supports the carbon-concentrating mechanism in Chlamydomonas reinhardtii. Plant Cell Physiol 51, 1453–1468 (2010).

22. Yamano, T., Toyokawa, C. & Fukuzawa, H. High-resolution suborganellar localization of Ca²⁺-binding protein CAS, a novel regulator of CO_2_-concentrating mechanism. Protoplasma 255, 1015–1022 (2018).

23. Shimamura, D. & Yamano, T. Regulatory logic of the Chlamydomonas CO_2_-concentrating mechanism: coupling carbon flux, energy supply, and pyrenoid architecture. Front Plant Sci 17, 1882169 (2026).

24. Fukuzawa, H. et al. Ccm1, a regulatory gene controlling the induction of a carbon-concentrating mechanism in Chlamydomonas reinhardtii by sensing CO_2_ availability. Proc Natl Acad Sci USA 98, 5347–5352 (2001).

25. Xiang, Y., Zhang, J. & Weeks, D. P. The Cia5 gene controls formation of the carbon-concentrating mechanism in Chlamydomonas reinhardtii. Proc Natl Acad Sci USA 98, 5341–5346 (2001).

26. Miura, K. et al. Expression profiling-based identification of CO_2_-responsive genes regulated by CCM1 controlling a carbon-concentrating mechanism in Chlamydomonas reinhardtii. Plant Physiol 135, 1595–1607 (2004).

27. Fang, W. et al. Transcriptome-wide changes in Chlamydomonas reinhardtii gene expression regulated by carbon dioxide and the CO_2_-concentrating mechanism regulator CIA5/CCM1. Plant Cell 24, 1876–1893 (2012).

28. Yoshioka, S. et al. The novel Myb transcription factor LCR1 regulates the CO_2_-responsive gene Cah1, encoding a periplasmic carbonic anhydrase in Chlamydomonas reinhardtii. Plant Cell 16, 1466–1477 (2004).

29. Chen, B. & Spalding, M. H. Investigation of the DNA binding ability of CIA5 in Chlamydomonas reinhardtii. Plant Mol Biol Rep 44, 42 (2026).

30. Wang, L. et al. Chloroplast-mediated regulation of CO_2_-concentrating mechanism by Ca²⁺-binding protein CAS in the green alga Chlamydomonas reinhardtii. Proc Natl Acad Sci USA 113, 12586–12591 (2016).

31. Shimamura, D. et al. A pyrenoid-localized protein SAGA1 is necessary for Ca^2+^-binding protein CAS-dependent expression of nuclear genes encoding inorganic carbon transporters in Chlamydomonas reinhardtii. Photosynth Res 156, 181–192 (2023).

32. Burlacot, A. et al. Alternative photosynthesis pathways drive the algal CO_2_-concentrating mechanism. Nature 605, 366–371 (2022).

33. Peltier, G. et al. Alternative electron pathways of photosynthesis power green algal CO_2_ capture. Plant Cell 36, 4132–4142 (2024).

34. Shimamura, D. et al. A nuclear CobW/WW-domain factor represses the CO_2_-concentrating mechanism in the green alga Chlamydomonas reinhardtii. Proc Natl Acad Sci USA 123, e2518136123 (2026).

35. Mackinder, L. C. M. et al. A spatial interactome reveals the protein organization of the algal CO_2_-concentrating mechanism. Cell 171, 133–147.e14 (2017).

36. Ramazanov, Z., Rawat, M., Henk, M. C., Mason, C. B., Matthews, S. W. & Moroney, J. V. The induction of the CO_2_-concentrating mechanism is correlated with the formation of the starch sheath around the pyrenoid of Chlamydomonas reinhardtii. Planta 195, 210– 216 (1994).

37. Sudol, M. et al. Characterization of the novel protein-binding module: the WW domain. FEBS Lett 369, 67–71 (1995).

38. Fei, C. et al. Modelling the pyrenoid-based CO_2_-concentrating mechanism provides insights into its operating principles and a roadmap for its engineering into crops. Nat Plants 8, 583–595 (2022).

39. Itakura, A. K. et al. A Rubisco-binding protein is required for normal pyrenoid number and starch sheath morphology in Chlamydomonas reinhardtii. Proc Natl Acad Sci USA 116, 18445–18454 (2019).

40. Young, T. R. et al. Calculating metalation in cells reveals CobW acquires CoII for vitamin B12 biosynthesis while related proteins prefer ZnII. Nat Commun 12, 1195 (2021).

41. Yamano, T. & Fukuzawa, H. Indirect immunofluorescence assay in Chlamydomonas reinhardtii. Bio-protocol 6, e1864 (2016).

42. Yamano, T, RNA-seq data. DDBJ Sequence Read Archive. https://ddbj.nig.ac.jp/search/entry/bioproject/ PRJDB43043. Deposited 30 August 2026.

43. Chlamydomonas Resource Center, Scientific and Community News. https://www.chlamycollection.org/. Accessed 30 August 2026.

44. US Department of Energy Joint Genome Institute, Chlamydomonas reinhardtii HCR1 gene (Cre16.g684650). Phytozome. https://phytozome-next.jgi.doe.gov/report/gene/Creinhardtii_v5_6/Cre16.g685050. Accessed 30 August 2026.

## SI References

1. D. Shimamura et al., A nuclear CobW/WW-domain factor represses the CO2-concentrating mechanism in the green alga Chlamydomonas reinhardtii. Proc. Natl. Acad. Sci. U.S.A. 123, e2518136123 (2026).

2. T. Yamano, C. Toyokawa, D. Shimamura, T. Matsuoka, H. Fukuzawa, CO2-dependent migration and relocation of LCIB, a pyrenoid-peripheral protein in Chlamydomonas reinhardtii. Plant Physiol. 188, 1081–1094 (2022).

3. D. Shimamura, T. Yamano, Regulatory logic of the Chlamydomonas CO2-concentrating mechanism: coupling carbon flux, energy supply, and pyrenoid architecture. Front. Plant Sci. 17, 1882169 (2026).

4. K. Katoh, D. M. Standley, MAFFT multiple sequence alignment software version 7: improvements in performance and usability. Mol. Biol. Evol. 30, 772–780 (2013).

5. S. Kumar, G. Stecher, M. Li, C. Knyaz, K. Tamura, MEGA X: Molecular Evolutionary Genetics Analysis across computing platforms. Mol. Biol. Evol. 35, 1547–1549 (2018).

6. J. P. Concordet, M. Haeussler, CRISPOR: intuitive guide selection for CRISPR/Cas9 genome editing experiments and screens. Nucleic Acids Res. 46, W242–W245 (2018).

7. J. G. Doench et al., Optimized sgRNA design to maximize activity and minimize off-target effects of CRISPR-Cas9. Nat. Biotechnol. 34, 184–191 (2016).

8. T. Yamano, H. Iguchi, H. Fukuzawa, Rapid transformation of Chlamydomonas reinhardtii without cell-wall removal. J. Biosci. Bioeng. 115, 691–694 (2013).

9. L. Wang et al., Chloroplast-mediated regulation of CO2-concentrating mechanism by Ca2+-binding protein CAS in the green alga Chlamydomonas reinhardtii. Proc. Natl. Acad. Sci. U.S.A. 113, 12586–12591 (2016).

10. T. Yamano et al., Light and low-CO2-dependent LCIB–LCIC complex localization in the chloroplast supports the carbon-concentrating mechanism in Chlamydomonas reinhardtii. Plant Cell Physiol. 51, 1453–1468 (2010).

11. A. Tachiki, H. Fukuzawa, S. Miyachi, Characterization of carbonic anhydrase isozyme CA2, which is the CAH2 gene product, in Chlamydomonas reinhardtii. Biosci. Biotechnol. Biochem. 56, 794–798 (1992).

12. T. Yamano et al., Characterization of cooperative bicarbonate uptake into chloroplast stroma in the green alga Chlamydomonas reinhardtii. Proc. Natl. Acad. Sci. U.S.A. 112, 7315–7320 (2015).

13. N. Ohnishi et al., Expression of a low CO2-inducible protein, LCI1, increases inorganic carbon uptake in the green alga Chlamydomonas reinhardtii. Plant Cell 22, 3105–3117 (2010).

14. C. Toyokawa, T. Yamano, H. Fukuzawa, Pyrenoid starch sheath is required for LCIB localization and the CO2-concentrating mechanism in green algae. Plant Physiol. 182, 1883–1893 (2020).

15. D. Kim et al., Graph-based genome alignment and genotyping with HISAT2 and HISAT-genotype. Nat. Biotechnol. 37, 907–915 (2019).

16. S. Anders, P. T. Pyl, W. Huber, HTSeq—a Python framework to work with high-throughput sequencing data. Bioinformatics 31, 166–169 (2015).

17. M. D. Robinson, D. J. McCarthy, G. K. Smyth, edgeR: a Bioconductor package for differential expression analysis of digital gene expression data. Bioinformatics 26, 139– 140 (2010).

18. T. Yamano, H. Fukuzawa, Indirect immunofluorescence assay in Chlamydomonas reinhardtii. Bio-protocol 6, e1864 (2016).

